# Multiple p38/JNK Mitogen-activated protein kinase (MAPK) signaling pathways mediate salt chemotaxis learning in *C. elegans*

**DOI:** 10.1101/2023.05.18.541291

**Authors:** Taoruo Huang, Kota Suzuki, Hirofumi Kunitomo, Masahiro Tomioka, Yuichi Iino

**Affiliations:** Department of Biological Sciences, Graduate School of Science, The University of Tokyo, Tokyo, Japan

## Abstract

Animals are able to adapt their behaviors to their environment. In order to achieve this, the nervous system plays integrative roles, such as perception of external signals, sensory processing, and behavioral regulations via various signal transduction pathways. Here genetic analyses of *C. elegans* found that mutants of components of JNK and p38 Mitogen-activated protein kinase (MAPK) signaling pathways, also known as stress-activated protein kinase (SAPK) signaling pathways, exhibit various types of defects in the learning of salt chemotaxis. *C. elegans* homologues of JNK MAPKKK and MAPKK, MLK-1 and MEK-1, respectively, are required to avoid salt concentrations experienced during starvation. In contrast, homologues of p38 MAPKKK and MAPKK, NSY-1 and SEK-1, respectively, are required for high-salt chemotaxis after conditioning. Genetic interaction analyses suggest that a JNK family MAPK, KGB-1, functions downstream of both signaling pathways to regulate salt chemotaxis learning. Furthermore, we found that the NSY-1/SEK-1 pathway functions in sensory neurons, ASH, ADF, and ASER, to regulate the learned high-salt chemotaxis. A neuropeptide, NLP-3, expressed in ASH, ADF, and ASER neurons, and a neuropeptide receptor, NPR-15, expressed in AIA interneurons that receive synaptic input from these sensory neurons, function in the same genetic pathway as NSY-1 / SEK-1 signaling. These findings suggest that this MAPK pathway may affect neuropeptide signaling between sensory neurons and interneurons, thus promoting high-salt chemotaxis after conditioning.

## Introduction

The ability to adapt is crucial in nature. In order to ensure their survival in their environment, organisms can sense, identify, and respond to ever-changing environmental stimuli. At the cellular level, the response to extracellular stimuli is an important feature of cellular regulation. Of the signaling pathways involved in such regulation, mitogen-activated protein kinase (MAPK) signaling pathways are a major subtype. Extracellular stimuli lead to sequential phosphorylation of each member of the MAP3K-MAP2K-MAPK cascade in the cytoplasm, which subsequently causes the alteration of transcription and translation in the nucleus (Hazzalin et al., 1997), thus achieving physiological regulation of the cell. Among the four groups of MAPKs, the JNK and p38 classes are activated by environmental stress stimuli such as ultraviolet light, heat, osmotic pressure, and pathogenic bacteria (Liu et al., 2002; Benndorf et al., 1994; Parker et al., 1998; Kim et al., 2002). JNK and p38 MAPKs are collectively called stress-activated protein kinases (SAPKs) and play important roles in response to stressful stimuli.

One strategy of living animals for adaptation to the environment to escape from an unpreferred place and move to a more comfortable one. Moreover, most animals have the innate ability to seek out optimal environmental conditions. The model experimental organism, *C. elegans*, can sense environmental stimuli, including odorants and water-soluble chemicals, such as salts, and learn these sensory stimuli to modulate their behavior (Bargmann, 2006). Additionally, worms are attracted to the salt concentration at which they have been fed, while they avoid this salt concentration if they have been starved (Saeki et al., 2001; Kunitomo et al., 2013). This form of associative learning, called salt chemotaxis learning, causes a switch between attraction and avoidance according to food availability and helps worms seek favorable habitats with abundant food. In salt chemotaxis learning, worms are stimulated by various environmental stimuli, including high salt concentrations, high osmotic pressures, and absence of food, all of which have a potential to activate SAPKs. Furthermore, conserved SAPK pathways in *C. elegans* are required for neural functions such as neuronal differentiation (Chuang and Bargmann, 2005), axon regeneration and degeneration (Nix et al., 2011; Ding et al., 2022), and forgetting of memory (Inoue et al., 2013). Therefore, we hypothesized that SAPKs might be involved in salt chemotaxis learning in *C. elegans*.

*C. elegans* has a relatively simple nervous system where all neurons have been identified individually, and connections among individual neurons have been well documented, which helps with understanding information processing within their neural circuits (White et al., 1986). Water-soluble chemicals are mainly detected by chemosensory neurons in the head, named amphid sensory neurons. Among them, ASE neurons are thought to primarily sense ambient salt concentrations and are crucial for salt chemotaxis (Bargmann and Horvitz, 1991). Neuron ablation experiments revealed that ASER, the right-side member of the left/right pair of ASE neurons, plays a dominant role in salt chemotaxis (Kunitomo et al., 2013). However, after ASE is killed, ADF, ASI, and ASG contribute to a residual response to several water-soluble molecules, including sodium and chloride ions (Bargmann and Horvitz 1991). The main nociceptor, ASH sensory neurons, also contribute to a balance between attraction and avoidance of salt, in concert with ASE, ASI, ADF, and perhaps ADL neurons (Hukema et al., 2006). These results suggested that salt chemotaxis learning could be controlled by several classes of sensory neurons.

Neuropeptides are generally used to transmit endogenous information and regulate various neuronal functions *in vivo*. For example, in *C. elegans*, neuropeptides are involved in the regulation of locomotion, reproduction, and the growth and differentiation of neurons. A previous study found that carboxypeptidase E, EGL-21, which is required for the production of mature neuropeptides, functions downstream of the insulin pathway to regulate salt chemotaxis learning (Nagashima et al., 2019), implicating the relationship between neuropeptide signaling and salt chemotaxis learning.

In this study, we show that in *C. elegans*, two conserved SAPK pathways are involved in salt chemotaxis learning. JNK signaling, carried by MLK-1(MAP3K) and MEK-1(MAP2K), mediates the avoidance of salt concentrations experienced during starvation. In contrast, p38 signaling, carried by NSY-1(MAP3K) and SEK-1(MAP2K), specifically mediates high-salt chemotaxis after conditioning under starvation with low salt and feeding with high salt. A MAPK from the JNK family, KGB-1, functions in both the JNK and p38 signaling pathways. Furthermore, we found that SEK-1 signaling functions in ASH, ADF, and ASER sensory neurons to mediate high-salt attractive learning under well-fed conditions, and neuropeptide signaling is likely involved in this regulation.

## Methods

### Strains and culturing

The *C. elegans* strains and transgenic lines used in this study are listed in Supplementary Table S1. Animals were incubated at 20℃ on 50 mM NaCl nematode growth medium (NGM) plates and seeded with fast-growing *E. coli* NA22 as a food source. Standard genetic methods were used to generate strains with double mutations by crossing (Brenner et al., 1974).

### DNA constructs and transgenesis

In order to generate *sek-1* and *grk-2* expression plasmids, we generated the destination vector carrying *sek-1(or grk-2)::sl2::cfp* cDNA (*sek-1* cDNA was a gift from Dr. Takeshi Ishihara), namely pDEST-SEK-1(or GRK-2)::SL2::CFP. Subsequently, promoter sequences were inserted upstream of *sek-1(or grk-2)::sl2::cfp* via the LR reaction with entry vectors carrying the promoter sequences (pENTR-promoter) using the GATEWAY cloning technique (Thermo Fisher Scientific, Inc., Waltham, MA, USA). The promoters used are listed in Supplementary Table S2. Next, cell-specific expression strains were generated by standard microinjection methods (Mello et al., 1991). The expression constructs for *sek-1* and *grk-2* were injected at 5 ng μl^-1^ along with the transformation marker (10 ng μl^-1^) and pPD49.26 (85 ng μl^-1^) as carrier DNA.

In order to generate a plasmid for the expression of guide RNA and Cas9, we added the *pmk-2* target sequence (atggggatgtcggcgaccat) or *mek-1* (cttggcatgggcagacctgg) to the pDD162 plasmid, including *eft-3p::Cas9* and empty *sgRNA* driven by the *U6* promoter (Dickinson et al., 2013), using inverse PCR. Each of these plasmid constructs was injected at 50 ng μl^-1^ along with a transformation marker *myo-3p::venus* (10 ng μl^-1^) and pPD49.26 (40 ng μl^-1^) as carrier DNA to isolate deletion mutants, *pmk-2(pe5707) IV.* and *mek-1(pe3745) X*.

### Salt Chemotaxis Learning Assay

To evaluate the learning of salt concentrations, in this study, a previously established salt chemotaxis learning assay was used (Kunitomo et al., 2013; Tomioka et al., 2016) (Supplementary Figure S1). Adult worms were washed with KP25 buffer (25 mM potassium phosphate (pH 6.0), 1 mM CaCl_2_, 1 mM MgSO_4_, 50 mM NaCl) and transferred to NGM plates with 25 or 100 mM NaCl in the absence or presence of food (NA22) for 6 h. After conditioning, the worms were placed in the center of a 9 cm test plate (25 mM potassium phosphate (PH 6.0), 1 mM CaCl_2_, 1 mM MgSO_4_, 2% agar) with a NaCl gradient of 35 to 95 mM and allowed to crawl for 45 min. Subsequently, 100*–*200 worms were used in each assay. The chemotaxis index was determined according to the following equation:

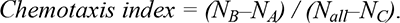

*N_A_* and *N_B_* are the numbers of worms in the low and high salt areas, respectively, while *N_all_* is the total number of worms on a test plate, and *N_C_* is the number of worms in the area around the starting position. The number *Nc* is not included in the calculation of chemotaxis index. The reason for omitting *Nc* is that the worms staying around the starting position can do so due to multiple reasons such as mobility defect. However, except for the *unc-43(e408)* with extremely severe mobility defect, we ensured that Nall-Nc was above 100 for each assay. An immobility index was separately calculated to represent the proportion of worms that stay in the C area:

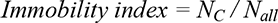

### Quantitative Real-Time PCR

The total RNA was extracted from wild-type animals and *sek-1(km4)* mutant animals before and after high-salt conditioning in the presence of food. cDNA was synthesized using the PrimeScript RT reagent kit (Takara) and used as templates for real-time PCR analysis using SYBR PreMix ExTaqII (Takara). The primers used are listed in Supplementary Table S3. The expression level was normalized to that of the wild-type before conditioning, and an *eef-1* gene was used as an internal standard.

### Statistical analyses

One-way ANOVA followed by Dunnett’s test and Bonferroni’s test was used to determine statistically significant differences between groups in salt chemotaxis assays. Additionally, for real-time PCR assays, Kruskal*–*Wallis and Mann*–*Whitney tests were used in statistical analyses. These analyses were performed using GraphPad Prism 9.4 software (GraphPad Software, La Jolla, CA).

## Results

### Mutants in the JNK and p38 pathways exhibited different defects in salt chemotaxis learning

Salt chemotaxis learning in *C. elegans* is subdivided into several different paradigms. The worms migrate to the concentrations of sodium chloride, which it has experienced under fed conditions. More specifically, the worms migrate to low (25 mM) and high (100 mM) salt concentrations after conditioning at low and high salt, respectively, under fed conditions; hereafter, these phenomena are called low-salt attractive and high-salt attractive learning. Conversely, *C. elegans* avoids the salt concentrations experienced under starvation conditions. They migrate to high and low salt concentrations after conditioning at low and high salt, respectively, under starved conditions (hereafter called low-salt avoidance and high-salt avoidance learning, respectively) (Supplementary Figure S1). In order to demonstrate the relevance of SAPK signaling pathways to salt chemotaxis learning, we carried out a salt chemotaxis learning assay using several mutants of the components of p38 MAPK or JNK pathways. We found that some mutants of the components of JNK and p38 pathways exhibited various types of salt chemotaxis learning defects. (All components are shown in Supplementary Figure S2) For the JNK pathway, deletion mutants of MLK-1(MAP3K) and MEK-1(MAP2K), *mlk-1(ok2471) and mek-1(pe3475)*, respectively, exhibited defects in avoidance learning of both high and low salt concentrations after starvation conditioning (Figure 1a). In contrast, a mutant of JKK-1 (MAP2K), *jkk-1(km2)*, showed normal salt chemotaxis learning (Supplementary Figure S3a). In the p38 MAPK pathway, missense mutants *nsy-1(ag3)* and the *sek-1(ag1)*, deletion mutant *sek-1(km4)*, and the double mutant *nsy-1(ag3); sek-1(km4)* exhibited defects specifically in high-salt chemotaxis after high salt/fed and low salt/starved conditioning (Figure 1b). Conversely, mutants of other p38 MAPK pathway components, DLK-1(MAP3K) and MKK-4 (MAP2K), showed normal salt chemotaxis learning (Supplementary Figure S3b). In order to test the genetic interaction between the JNK and p38 pathways, we constructed the *sek-1(ag1); mlk-1(ok2471)* double mutant. The *sek-1* mutation further decreased high-salt migration after high-salt attractive learning in the *mlk-1* mutant background. Furthermore, the *sek-1* mutation promoted low salt migration after high-salt avoidance learning; therefore, it appeared to suppress the high-salt avoidance defect in the *mlk-1(ok2471)* mutant (Figure 1a). This additive phenotype in the double mutant indicated that they function in parallel pathways to mediate salt chemotaxis learning.

**Figure 1.**
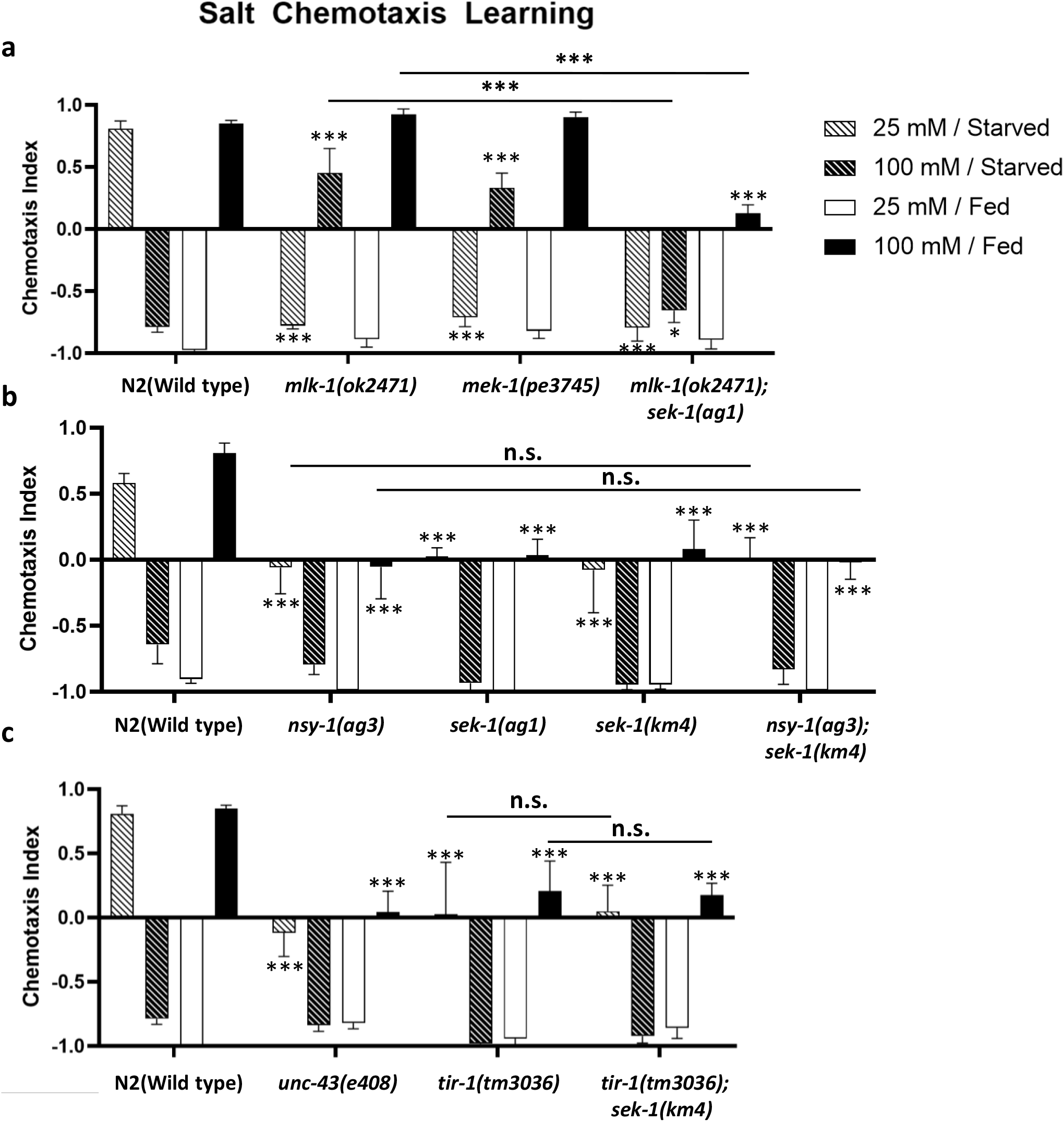
Mutants of the *mlk-1-mek-1* JNK and *nsy-1*-*sek-1* p38 MAPK pathways exhibit salt chemotaxis learning defects. Salt chemotaxis learning in the wild-type *C. elegans* and mutants of the components of JNK and p38 MAPK pathways. The chemotaxis index represents the direction and extent of salt chemotaxis. When this index is positive and negative, *C. elegans* shows high and low salt migration, respectively. (a) Mutants of *mlk-1* and *mek-1* exhibit salt avoidance learning defects under starved conditions. The *mlk-1; sek-1* double mutant shows the additive phenotype of those in the *mlk-1* and *sek-1* single mutants: The double mutant exhibits strong defects in low-salt avoidance and high-salt attractive learning (25 mM/starved and 100 mM/fed) with a weak defect in high-salt avoidance learning (100 mM/starved). (b) Mutants of *nsy-1* and *sek-1* exhibit defects in high-salt attraction after conditioning with 25 mM / starved and 100 mM / fed conditioning. The *nsy-1; sek-1* double mutant shows salt chemotaxis learning as defective as the single mutants. (c) Mutants of *unc-43* and *tir-1* exhibit defects in low-salt avoidance and high-salt attractive learning. The *tir-1; sek-1* double mutant shows salt chemotaxis learning as defective as the single mutants. The bars and error bars represent the mean values and SEM, respectively. ANOVA with Dunnett’s post hoc test (vs wild-type or single mutant (*mlk-1; nsy-1; tir-1*)): *P < 0.05, ***P < 0.001, n.s. not significant. N = 9–12 assays.

The ortholog of human SARM, TIR-1, is an activator of NSY-1 (Couillault et al., 2004). Moreover, the calcium-calmodulin-dependent protein kinase II (CaMKII) ortholog, UNC-43, reportedly binds to TIR-1 to recruit NSY-1 to presynaptic sites in AWC sensory neurons (Chuang and Bargmann 2005). We performed a salt chemotaxis learning assay using *tir-1(tm3036)* and *unc-43(e408)* mutants, which exhibited defects in high-salt chemotaxis similar to the *nsy-1* and *sek-1* mutants. This suggests that TIR-1 and UNC-43 are also required for salt chemotaxis learning (Figure 1c). Furthermore, the *sek-1(km4)* mutation did not further decrease high-salt chemotaxis in *tir-1(tm3036)* and *nsy-1(ag3)* mutant backgrounds (Figure 1b, c), indicating that *tir-1*, *nsy-1*, and *sek-1* function in the same genetic pathway, similar to those in stress responses and AWC development, to regulate salt chemotaxis learning.

Although substantial defects were observed in MAPKKK and MAPKK mutants of the JNK and p38 pathways, most mutants of known p38 and JNK MAPKs exhibited normal salt chemotaxis learning. In mutants of the MAPKs of the p38 family, the deletion mutants *pmk-1(km25)*, *pmk-2(pe5707),* and *pmk-3(ok169)* exhibited no obvious defect in salt chemotaxis learning. (Figure 2a) In MAPK mutants of the JNK family MAPKs, *jnk-1(gk7)* and *kgb-2(gk361)* exhibited no obvious defect (Figure 2b), whereas only *kgb-1(um3)* exhibited strong defects in low-salt avoidance and high-salt attractive learning (Figure 2c) reminiscent of the salt chemotaxis phenotype of *mlk-1(ok2471); sek-1(ag1)* mutants. In order to test the genetic interactions between both pathways (*mlk-1-mek-1* and *nsy-1-sek-1*) and *kgb-1*, we generated double mutants, *mlk-1(ok2471); kgb-1(um3)* and *kgb-1(um3); sek-1(km4).* Both the double mutant *mlk-1; kgb-1* and the double mutant *kgb-1; sek-1* exhibited chemotaxis defects as severe as those in the single mutant *kgb-1(um3)* (Figure 2c). These results suggest that *kgb-1* functions in the same pathway as *sek-1* and *mlk-1* and may act in the MLK-1-MEK-1 and NSY-1-SEK-1 cascades to mediate salt chemotaxis learning (Figure 3).

**Figure 2.**
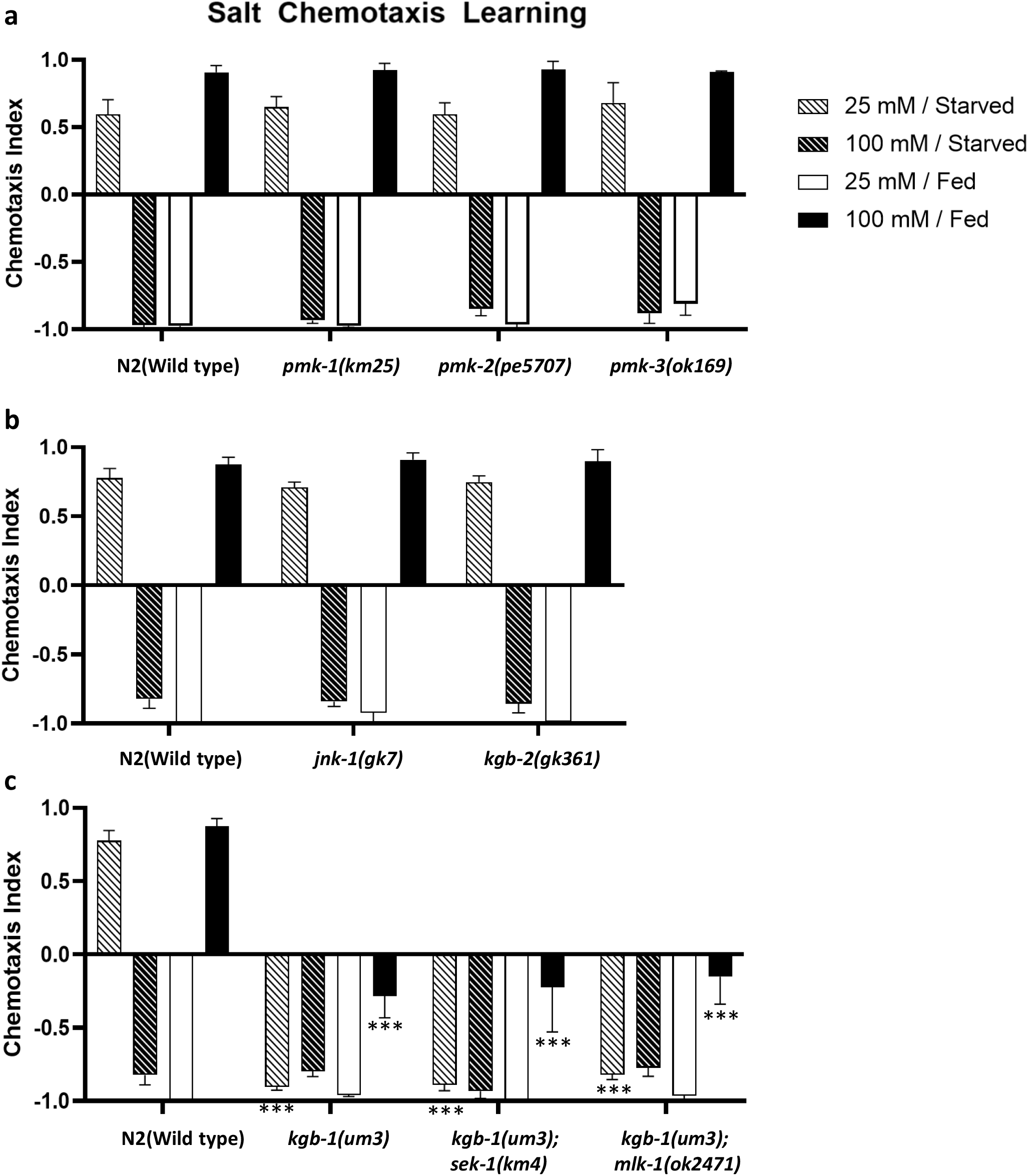
The *kgb-1* mutant exhibits salt chemotaxis learning defects. Mutants of p38 MAPKs, *pmk-1*, *pmk-2*, *pmk-3* (a) and JNKs, *kgb-2*, *jnk-1* (b) exhibit normal salt chemotaxis learning. (c) A *kgb-1* mutant exhibits defects in high-salt chemotaxis after low-salt avoidance and high-salt attraction learning. Both *kgb-1; sek-1,* and *kgb-1; mlk-1* double mutant animals exhibit the same defects as the *kgb-1* single mutant animals. The bars and error bars represent the mean values and SEM, respectively. ANOVA with Dunnett’s post hoc test (vs wild-type or *kgb-1*): ***P < 0.001, n.s. not significant. N = 9–12 assays.

**Figure 3.**
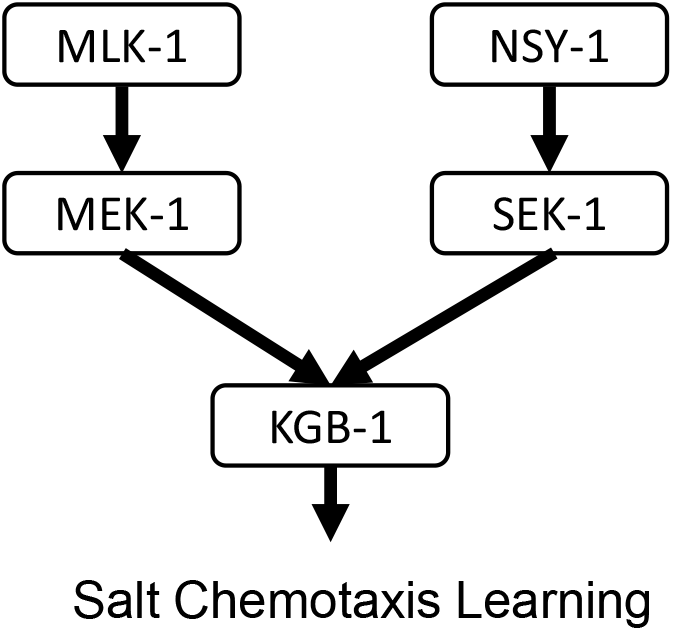
Summary of putative p38/JNK MAPK cascades in salt chemotaxis learning. The NSY-1(MAP3K)-SEK-1(MAP2K) pathway specifically mediates high-salt migration, and the MLK-1(MAP3K)-MEK-1(MAP2K) pathway mediates low and high-salt avoidance learning. KGB-1 (MAPK) presumably functions downstream of both MAP3K-MAP2K pathways to mediate salt chemotaxis learning.

### SEK-1 signaling redundantly regulates high-salt chemotaxis learning in multiple sensory neurons

In order to determine where the SEK-1 signaling pathway functions, cell-type-specific rescue experiments were carried out with *sek-1* cDNA. First, we performed rescue experiments using the *rgef-1* and *ges-1* promoters to test if SEK-1 functions in neurons and the intestine, respectively. Pan-neuronal *sek-1* expression fully rescued high-salt attractive learning in a well-fed condition, while *sek-1* expression in the intestine had no effect (Figure 4a). Conversely, *sek-1* expression in either the nervous system or the intestine did not rescue low-salt avoidance learning under starvation conditions (Figure 4b), implying that SEK-1 may differentially regulate high-salt migration between food-associated attractive and starvation-associated avoidance learning.

**Figure 4.**
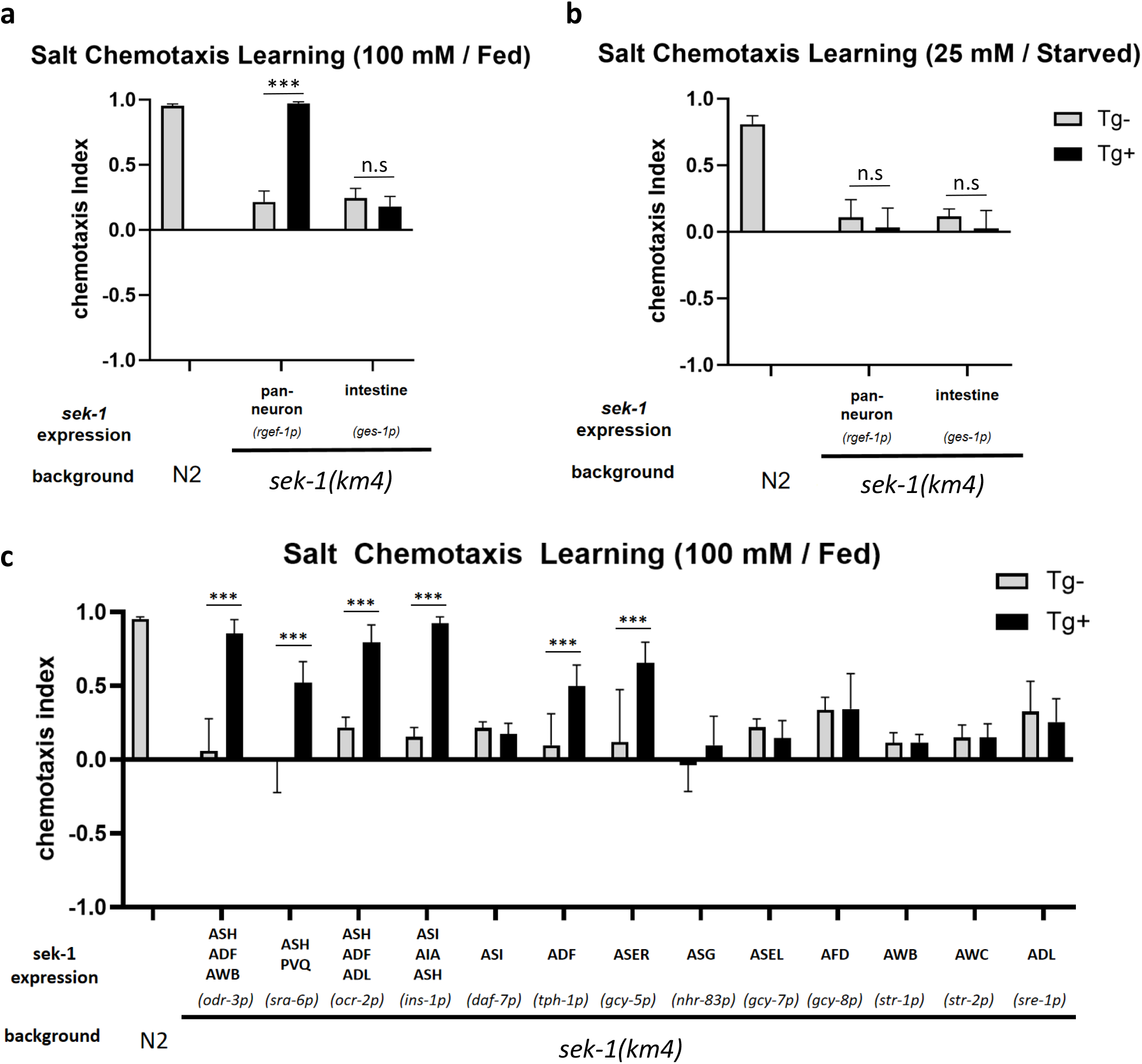
SEK-1 functions in sensory neurons ASER, ASH, and ADF in high-salt attractive learning. Cell-specific rescue experiments of salt chemotaxis learning defects in the *sek-1* mutant. Cells or tissues exhibiting promoter-driven expression are indicated, along with the name of the promoter in parentheses. The high-salt attractive learning defect was rescued by *sek-1* expression in neurons but not in the intestine (a), while the low-salt avoidance learning defect could not be rescued by *sek-1* expression in neurons or the intestine (b). The *odr-3, sra-6, ocr-2, ins-1, tph-1,* or *gcy-5* promoter, which drives expression in restricted cell classes as indicated, significantly rescues the high-salt attractive learning defect (c). Tg: transgene. The bars and error bars represent the mean values and SEM, respectively. ANOVA with Bonferroni’s post hoc test (Tg- vs Tg+): ***P < 0.001, n.s. not significant. N = 12 assays.

The SEK-1 signaling has been shown to play roles in various neuronal processes, including the development of AWC olfactory neurons, forgetting of olfactory memory by acting in AWC, and the biosynthesis of serotonin in ADF chemosensory neurons in response to pathogenic stimuli. Thus, we focused on sensory neurons as candidate functional sites of *sek-1* in salt chemotaxis learning (Sagasti et al., 2001; Inoue et al., 2013; Shivers et al., 2009). By using neuron-type-specific promoters, *sek-1* was expressed in several chemosensory neurons. The expression of *sek-1* only in the ASER neuron partially rescued the defect of the *sek-1* mutant in high-salt attractive learning under well-fed conditions. The expression of *sek-1* by the *odr-3* (ASH, ADF, and AWB selective) or *ocr-2 (*ASH, ADF, and ADL selective) promoters fully rescued the defect in the *sek-1* mutant. Among these neurons, *sek-1* expression only in ASH or ADF but not in ADL or AWB rescued the defect in the *sek-1* mutant (Figure 4c). These results suggest that *sek-1* expression in ASH, ADF, or ASER neurons is sufficient for high-salt attractive learning. However, in low-salt avoidance learning, *sek-1* expression in any neuron we tested, including some interneurons, did not rescue the defect in the *sek-1* mutant, except that *sek-1* expression in ASH by the *sra-6* promoter slightly rescued the defect (Supplementary Figure S4a). The *sra-6* promoter has been reported to be expressed mainly in ASH, PVQ interneurons, and the intestine. Although expression using other promoters was carried out to mimic the *sra-6* promoter-driven expression pattern, no rescue was observed (Supplementary Figure S4b), implying that SEK-1 signaling mainly acts at a site different from the neurons and the intestine and/or the precise expression level of SEK-1 is important for the rescue of the defects in low-salt avoidance.

### GRK-2 functions in the SEK-1 pathway in salt chemotaxis learning

In salt chemotaxis, a G protein-coupled receptor kinase (GRK), GRK-2, has been proposed to function in ASH sensory neurons to promote salt attraction, and *grk-2* mutants exhibited severely impaired chemotaxis and increased avoidance of salt (Hukema et al., 2006). Next, we examined a possible interaction between GRK-2 and the p38 pathway in regulating salt chemotaxis learning. GRKs are a family of protein kinases named for their abilities to modulate G protein-coupled receptor (GPCR) activities. However, in mammals, GRKs can also transmit signals via the phosphorylation of non-GPCR proteins, including p38 and JNKs. In human cells *in vitro*, GRK2 modulates p38-dependent physiological processes such as LPS-dependent TNF alpha secretion by regulating the activation of p38 (Peregrin et al., 2006), and GRK3 is required for JNK activation (Kuhar et al., 2015). In *C. elegans*, there are two GRK homologs, GRK-1 and GRK-2. The *C. elegans* ortholog of mammalian GRK2/3, GRK-2, but not GRK-1, is involved in SAPK-dependent stress responses, such as oxidative and heat stress (Henry et al., 2021; Heilman et al., 2020). These previous reports implied that GRK-2 may modulate SAPKs in *C. elegans* and be involved in salt chemotaxis learning regulation. Furthermore, beta-arrestin in mammals is known to cooperate with GRK in GPCR desensitization and promote p38 MAPK phosphorylation in mammals (Vroon et al., 2006; Bruchas et al., 2006). In the *C. elegans* genome, there is a single beta-arrestin coding gene, *arr-1*, which is required for adaptation to various odorants (Palmitessa et al., 2005) and may also function in salt chemotaxis learning. Subsequently, to examine the requirements of GRK-2 and ARR-1 in salt chemotaxis learning, we tested salt chemotaxis learning using a deletion mutant *grk-2(gk268)*, a missense mutant *grk-2(rt97)*, and a deletion mutant *arr-1(ok401)*. Both alleles of *grk-2* exhibited defects in high-salt attractive and low-salt avoidance learning, while *arr-1(ok401)* exhibited no defects in salt chemotaxis learning (Figure 5a), which implies that GRK-2 mediates high-salt chemotaxis learning without the requirement of beta-arrestin. Notably, although the defect in high-salt attractive learning in *grk-2* mutants appears to be at the same level compared to *sek-1* mutants, the defect in low-salt avoidance learning of *grk-2* mutants is much more severe. Furthermore, we generated the *grk-2(gk268); sek-1(km4)* double mutant and tested salt chemotaxis learning to determine whether *grk-2* functions in the same genetic pathway as *sek-1*. Thus, we observed that the double mutant exhibited defective salt chemotaxis at the same level as the *grk-2* single mutant after high-salt attractive learning (Figure 5a). The absence of additive phenotypes by the *sek-1* mutation under the *grk-2* mutant suggested that *sek-1* and *grk-2* function in the same genetic pathway to mediate high-salt attractive learning. On the other hand, low-salt avoidance defects in *grk-2* and *grk-2; sek-1* mutants are more severe than the *sek-1* mutant, suggesting that GRK-2 may regulate low-salt avoidance learning via SEK-1 and other signaling mechanisms.

**Figure 5.**
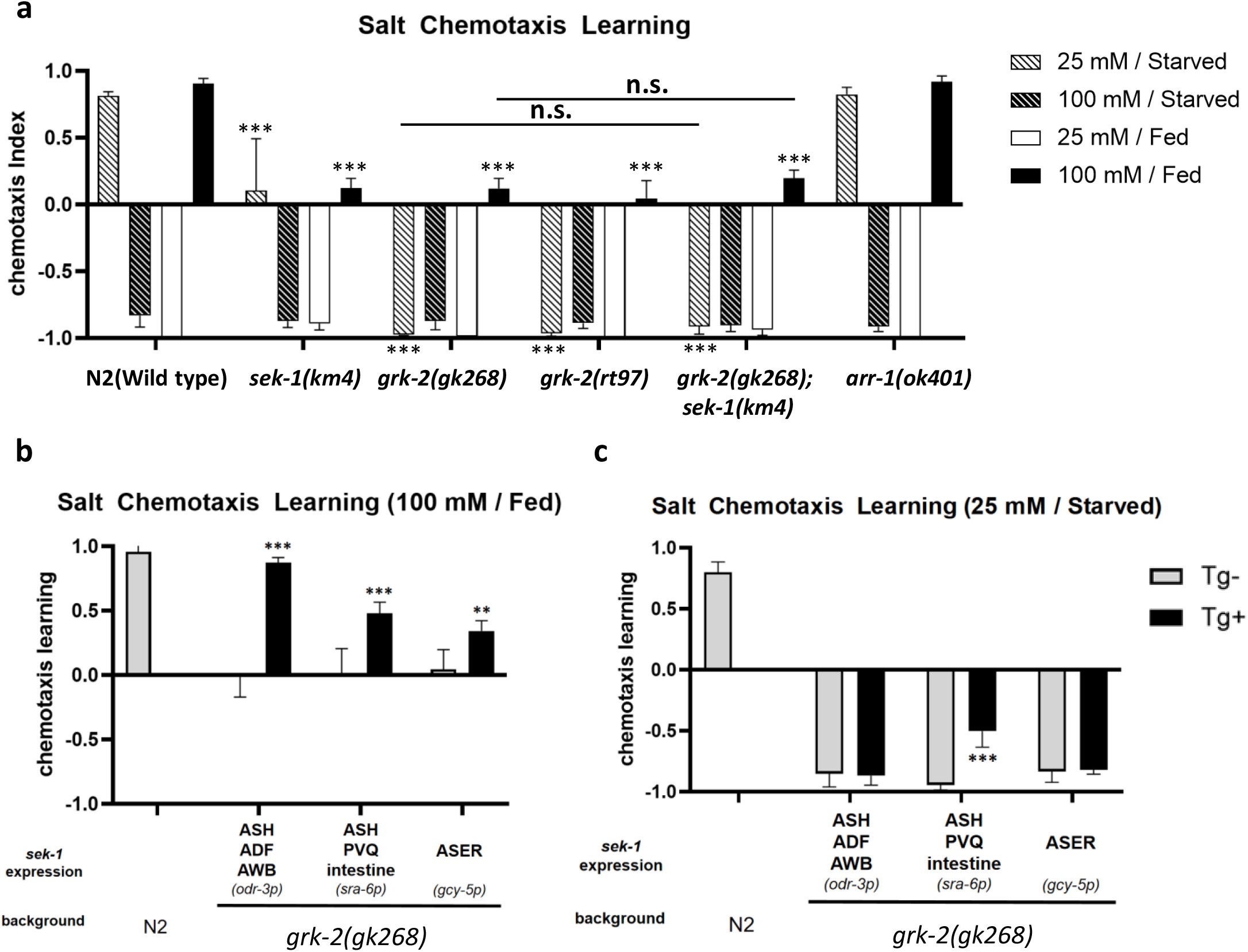
GRK-2 functions in high-salt attractive learning in the same genetic pathway as SEK-1. Salt chemotaxis assays were performed in wild-type animals, mutants, and transgenic *grk-2* mutants. Cells or tissues in which the promoter drives expression is indicated, along with the name of the promoter in parentheses. (a) *grk-2* mutant animals exhibit defects in both low-salt avoidance and high-salt attractive learning, while the *arr-1* mutant exhibits normal salt chemotaxis learning. The defects in the *grk-2* mutant are much more severe than in the *sek-1* mutant. The *grk-2; sek-1* double mutant is as defective as the *grk-2* single mutant in salt chemotaxis learning. ANOVA with Dunnett’s post hoc test (vs wild-type or *grk-2*): ***P < 0.001, n.s. not significant. N = 9 assays. (b) Similar to the rescue experiment in *sek-1* mutants (Fig. 3), *grk-2* expression by the *odr-3*, *sra-6* or *gcy-5* promoters restored high-salt attractive learning in the *grk-2* mutant. (c) Low-salt avoidance learning is rescued only by expressing *grk-2* using the *sra-6* promoter. The bars and error bars represent the mean values and SEM, respectively. ANOVA with Bonferroni’s post hoc test (Tg- vs Tg+): ***P < 0.001. N = 9 assays.

Next, cell-type-specific rescue experiments were performed to determine whether GRK-2 functions in sensory neurons where SEK-1 functions to mediate salt chemotaxis learning. We used the *odr-3*, *sra-6*, and *gcy-5* promoters for GRK-2 expression. As expected, using the *odr-3* or *gcy-5* promoter, the attractive learning defect, but not the avoidance learning defect in *sek-1* mutant animals, could be rescued, and the expression by the *sra-6* promoter restored both attractive and avoidance learning (Figure 5b, c). In conclusion, *grk-2* functions in the same genetic pathway as *sek-1* and contributes to regulating salt chemotaxis learning in sensory neurons.

### NPR-15 promotes high-salt chemotaxis in the same pathway as SEK-1 signaling

Although the NSY-1-SEK-1 pathway plays an important role in cell response and cell fate determination, the regulatory mechanism downstream of this pathway is not yet clear. According to mammalian studies, the substrates of SAPKs include several transcription factors and protein kinases. Through these direct targets, SAPKs achieve transcriptional (Hazzalin et al., 1997; Huttunen et al., 1998) and post-transcriptional regulation (Lee et al., 1994; Kontoyiannis et al., 1999; Rutault et al., 2001). Furthermore, some studies have shown that JNK and p38 MAPKs cooperatively regulate the synthesis and release of cytokines (Oltmanns et al., 2003). In *C. elegans*, the SEK-1 p38 signaling pathway promotes the expression of antimicrobial peptides (AMPs), such as NLP-29, in response to fungal infection, injury, and aging (E et al., 2018). In light of these findings, we examined downstream regulation of SEK-1 signaling in salt chemotaxis learning. Here, we tested neuropeptides abundantly expressed in ASER, ASH, and ADF to demonstrate their contribution to salt chemotaxis learning. Based on the single-cell transcriptome sequencing database (CeNGEN), we focused on *ins-27*, *nlp-3*, *nlp-14*, and *nlp-47* neuropeptides, which are expressed in all three types of sensory neurons. We found that *nlp-3(tm3023)* mutants exhibited a slight defect in high-salt attractive learning, while *nlp-47(tm13464)* mutant animals exhibited a severe defect throughout chemotaxis learning (Figure 6a). In order to test the genetic interaction between *nlp-3* and *sek-1*, we constructed the *nlp-3; sek-1* double mutant. The *nlp-3* mutant did not further decrease high-salt migration after high-salt attractive learning in the *sek-1* mutant background, -suggesting a possibility that *nlp-3* and *sek-1* may function in the same genetic pathway to mediate high-salt attractive learning. Given the relatively mild phenotype of the *nlp-3(tm3023)* mutants, we also tested a *nlp-3* overexpression strain. The overexpression strain exhibited a higher chemotaxis index under all chemotaxis learning conditions (Figure 6c), consistent with the notion that *nlp-3* promotes high-salt attraction. The result also suggested that expression level of *nlp-3* can critically affect the behavior. We therefore tested the expression levels of *nlp-3* with or without conditioning in the wild-type and the *sek-1* mutant. As a result, quantitative PCR analysis showed that the *nlp-3* mRNA level did not change after high salt/fed conditioning and by the *sek-1* mutation (Supplementary Figure S5).

**Figure 6.**
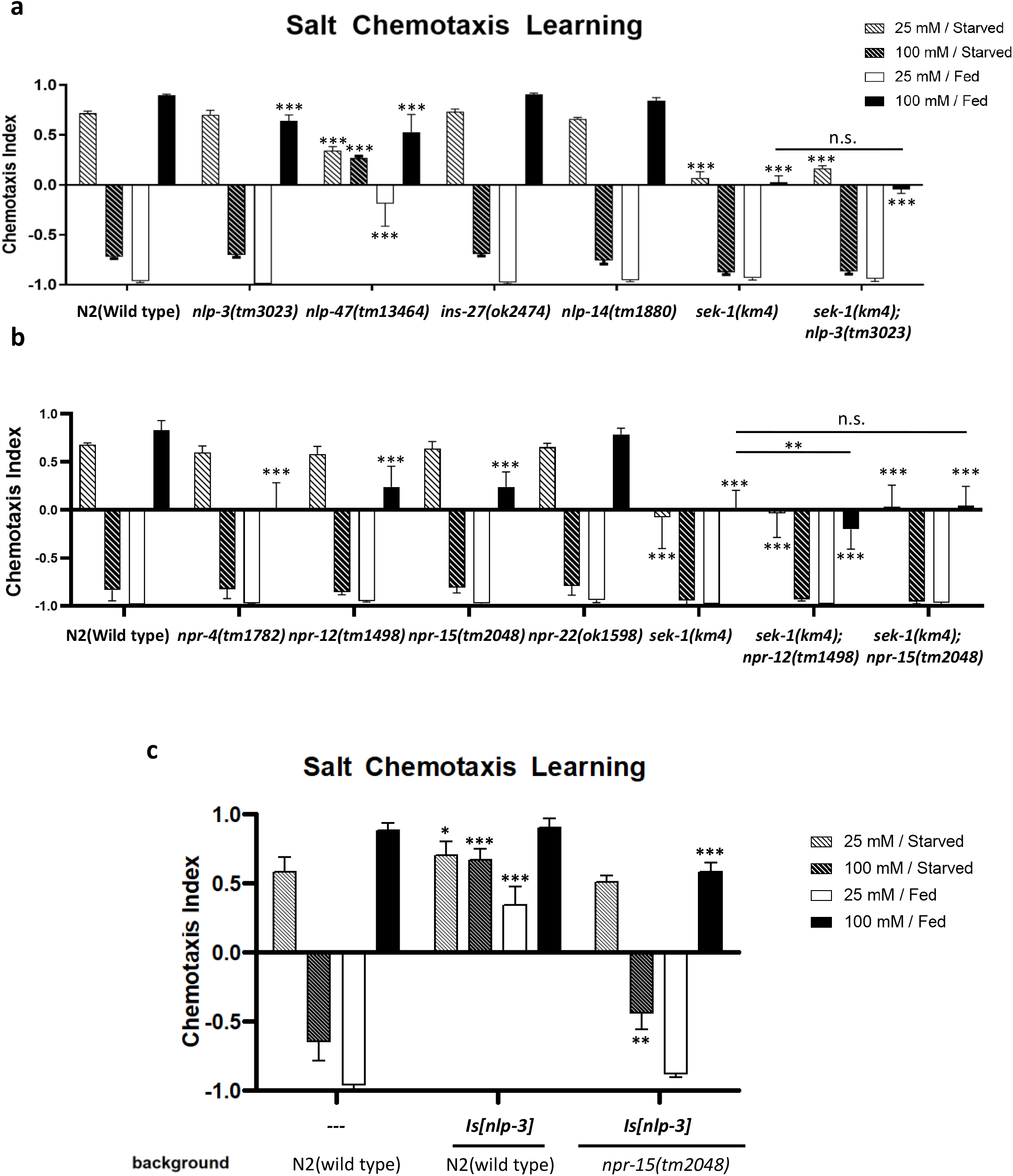
NLP-3 and NPR-15 promote high-salt attractive learning on the same pathway as SEK-1. Salt chemotaxis learning in mutants of neuropeptides expressed in all of ASER, ASH, and ADF sensory neurons (a) and neuropeptide receptors abundantly expressed in AIA interneurons (b). (a) The *nlp-3* mutants exhibited a high-salt attractive learning defect, and the *nlp-47* mutants exhibited a severe defect in all paradigms of salt chemotaxis learning. Additionally, the double mutant *nlp-3; sek-1* exhibits the high-salt attractive learning defect, which is statistically no different from that in the *sek-1* single mutant. (b) The *npr-4*, *npr-12*, and *npr-15* mutants exhibit high-salt attractive learning defects, also, the *npr-15; sek-1* double mutant exhibits the high-salt attractive learning defect, which shows no statistical difference to those in the *sek-1* single mutant. The bars and error bars represent the mean values and SEM, respectively. ANOVA with Dunnett’s post hoc test (vs wild-type or *sek-1*): **P < 0.01, ***P < 0.001, n.s. not significant. N = 12–15 assays. (c) Worms overexpressing *nlp-3* by its own promoter exhibit a higher salt chemotaxis index compared to the wild-type, and this effect was suppressed by the *npr-15* mutation. The bars and error bars represent the mean values and SEM, respectively. ANOVA with Dunnett’s post hoc test (vs wild-type). *P < 0.05, **P<0.01, ***P < 0.001. N = 6-12 assays.

A previous study has shown that AIA, AIB, and AIY, which are interneurons downstream of ASER, play an important role in salt chemotaxis learning (Mabardi et al., 2023). This study revealed that the ablation of AIA leads to a defect in high-salt attractive learning, while the collective ablation of AIA with AIB or AIY leads to more severe defects in high-salt attractive learning. Moreover, single ablation of AIB or AIY has no substantial effect on high-salt attractive learning (Mabardi et al., 2023). AIA ablation also leads to severe defects in high-salt avoidance learning in a different salt chemotaxis assay format (Tomioka et al., 2006). Based on the connectome database (White et al., 1986), ASH and ADF neurons also send synaptic inputs to AIB and AIY, respectively, and both ASH and ADF send inputs to AIA (Supplementary Figure S6).

Therefore, we focused on AIA neurons and searched for neuropeptide receptors expressed in AIA using the CeNGEN database. We hypothesized that neuropeptide receptors expressed in AIA might receive input from ASER, ASH, and ADF neurons via neuropeptides, such as NLP-3, to regulate salt chemotaxis learning. We tested *npr-15* and *npr-12,* expressed in AIA and AIZ, *npr-4,* expressed in AIA, AIB, AIY, AIZ, and *npr-22,* expressed only in AIA. Among these neuropeptide receptors, an ortholog of the human angiotensin II receptor (AGTR), *npr-15*, required for blood pressure and sodium retention, shows a relatively high expression in AIA. As a result, *npr-15*, *npr-12*, and *npr-4* mutants exhibited defects in high-salt attractive learning (Figure 6b). We constructed *npr-15; sek-1* double mutants and *npr-12; sek-1* double mutants. *npr-15; sek-1* double mutant animals exhibited the same defect as *sek-1* single mutant animals, while the *npr-12; sek-1* double mutant animals exhibited a more severe defect than the *sek-1* single mutant animals (Figure 6b), indicating that *npr-15* functions in the same genetic pathway as *sek-1*. To further investigate the possibility that NPR-15 is a receptor for NLP-3, *nlp-3* was overexpressed in *npr-15* mutant animals to observe whether the overexpression phenotype of *nlp-3* is suppressed by the mutation of *npr-15*. As a result, in contrast to overexpression in the wild-type background, overexpressing *nlp-3* in *npr-15(tm2048)* did not cause a significant change in salt chemotaxis learning <u>(Figure 6c)</u>, suggesting that the promotion effect of *nlp-3* on high-salt attraction requires *npr-15*, which implies that NPR-15 may be a receptor for NLP-3. These results suggest that *sek-1* might regulate high-salt attractive learning via neuropeptide signaling.

## Discussion

The ability of animals to choose their own habitat constitutes a central part of their environmental adaptability. In order to achieve this, animals perceive external information and process the information according to past and current environmental signals. In this study, we reveal that two conserved SAPK pathways, MLK-1-MEK-1 and NSY-1-SEK-1, function in parallel to regulate salt chemotaxis learning. MLK-1-MEK-1 mutants exhibited low- and high-salt avoidance learning defects, while NSY-1-SEK-1 mutants exhibited low-salt avoidance, and high-salt attractive learning defects. Furthermore, *mlk-1* and *sek-1* double mutants exhibited defects in both low-salt avoidance and high-salt attractive learning, but the high-salt avoidance learning defect is suppressed in double mutants; this phenotype is similar to that of the *kgb-1* mutant. By conducting genetic interaction analyses, we show that KGB-1 functions in the same genetical pathway as MLK-1 and SEK-1, implying that KGB-1 regulates both avoidance and attractive learning associated with salt and food availability under the coordinated control of the MLK-1 and SEK-1 signaling pathways (Figure 3). As previously mentioned, the NSY-1-SEK-1 signaling pathway plays multiple roles in biological processes, including mediating innate immunity in response to bacterial infection, neurosecretory control of serotonin, and asymmetry generation in neural development. PMK-1, a p38 MAPK, functions as a substrate of SEK-1 during the regulation of pathogen defense in peripheral tissues. However, in the nervous system, the substrate of SEK-1 appears to be divergent. In the regulation of the determination of AWC asymmetric cell fate and pathogen-induced serotonin biosynthesis in ADF, PMK-2 functions redundantly with PMK-1 downstream of SEK-1 (Pagano et al., 2015). Conversely, a JNK MAPK, JNK-1, mediates the forgetting of olfactory memory downstream of SEK-1 signaling in AWC neurons (Inoue et al., 2013). In this study, we propose that KGB-1, a member of the JNK family, contributes to the salt chemotaxis learning downstream of NSY-1-SEK-1 signaling. These results imply that several kinds of signaling acts downstream of SEK-1 in neurons to regulate various neural functions. Moreover, KGB-1 functions downstream of MLK-1-MEK-1 JNK signaling in neuronal and behavioral regulations, including synapse formation, axon regeneration, and pathogen avoidance (Baker et al., 2015; Nix et al., 2011, Melo and Ruvkun, 2012). In summary, multiple combinations of crosstalk between the p38 and JNK MAPK pathways are involved in controlling the development and functions of the nervous system. It is worth noting that mutants of *grk-2* showed a severe salt chemotaxis learning defect similar to the *mlk-1; sek-1* double and *kgb-1* single mutants. Based on the cell-specific rescue experiments and the genetic interaction analysis between mutations of *grk-2* and *sek-1*, we showed that *grk-2* and *sek-1* function in the same pathway in chemosensory neurons. Taken together with the report that mammalian GRK regulates both p38 and JNK signaling pathways, it is possible that GRK-2 regulates KGB-1 via both the NSY-1-SEK-1 p38 and MLK-1-MEK-1 JNK pathways in salt chemotaxis learning. To verify this hypothesis, it will be needed to identify the functional site of the MLK-1-MEK-1 pathway and its relationship with GRK-2.

Besides salt chemotaxis learning defects, some mutants also exhibited defects in mobility during the salt chemotaxis assay, manifested as the proportion of animals staying at the start point, which was significantly increased compared with wild type animals. *C. elegans* moves using a sinusoidal pattern of body bends, and the movement is generated by a complex interplay of muscles and neurons. In fact, it has been reported that mutants of *nsy-1* or *sek-1* cause low rate of body bends and amplitude (Hoyt et al., 2017), which might contribute to the immobility phenotype. Here, we found that mutants of the *mlk-1-mek-1* JNK and *nsy-1-sek-1* p38 MAPK pathways all exhibit a relatively higher immobility index (Supplementary Table S4). Interestingly, while expressing *sek-1* in neurons restored high-salt attractive learning, the immobility phenotype was not significantly rescued, suggesting SEK-1 may regulate mobility in non-neuronal sites.

ASE neurons are a crucial regulator of salt chemotaxis (Bargmann and Horvitz, 1991), and the functional asymmetry of left- and right-sided ASE neurons are largely reflected in the regulation of salt chemotaxis learning. The left-sided ASEL neuron is required for the attraction to sodium ions, whereas the right-sided ASER neuron is required for low- and high-salt attractive learning (Kunitomo et al., 2013; Wang et al., 2017). The *dyf-11* mutant animals are altered in the cilium biogenesis of sensory neurons (Kunitomo and Iino, 2008), thus causing severe defects in salt sensing and salt chemotaxis learning. The salt attractive learning was fully restored by expressing *dyf-11* cDNA in the ASER neuron, which allows normal ciliogenesis in ASER (Kunitomo et al., 2013), i.e., sensory input of ambient salt information to only the ASER neuron is sufficient for salt chemotaxis learning. In this study, we show that SEK-1 signaling in ASER promotes high-salt attractive learning but has no effect when it is expressed in ASEL, further supporting the importance of functional asymmetry of ASE neurons. Additionally, SEK-1 signaling in ASH and ADF also significantly restored the high-salt attractive learning defect in the *sek-1* mutant. GRK-2 expressed in ASH restored the high-salt attractive learning in the *grk-2* mutant. Therefore, normal functions of ASH and ADF are also required for high-salt attractive learning. In a previous report, researchers suggested that ADF may contribute to a residual response to the salt attraction after ASE is killed (Bargmann and Horvitz 1991). Additionally, ASH neurons have been reported to function along with ASE in salt chemotaxis plasticity after exposure to a high-salt containing buffer (Hukema et al., 2006; Dekkers et al., 2021). These findings are consistent with the assertion that multiple chemosensory neurons, including ADF and ASH, support the function of ASER in high-salt migration. The functions of ADF neurons have been reported to respond to stresses from pathogenic bacteria, which are mediated by p38 MAPK pathway activity (Shivers et al., 2009). ASH neurons receive multiple stress signals, including high osmolality and copper ions (Hilliard et al., 2005). It will be important to determine whether NSY-1-SEK-1 p38 MAPK signaling affects ADF and ASH functions in high-salt attractive learning. Another major remaining question is whether these mutants simply avoid high salt, regardless of learning, or have a defect in learning of several salt concentrations. In addition to conditioning with high (100 mM) or low (25 mM) salt, we tested salt chemotaxis after salt conditioning at 50 mM, a concentration around the center region in the chemotaxis test plate. As a result, wild type animals showed neither high nor low salt migration bias, while the mutant strains exhibited lower chemotaxis index (Supplementary Figure S7). Although the mutant worms appear to move lower salt regardless of salt concentrations during conditioning, we cannot still rule out the possibility that the mutant worms have defects learning of the medium salt concentration. Moreover, based on the results of the neuron-specific rescue experiment, in which neuronal *sek-1* expression restored the chemotaxis after high-salt attractive learning but not after low-salt avoidance learning, neuronal SEK-1 signaling may play a role in learning or learned chemotaxis in specific conditions, such as in the presence of food. Further study is needed to determine which process of salt chemotaxis learning the p38 MAPK pathway is involved, which is an important step toward better understanding of the functions of p38 MAPK pathway in behavioral plasticity.

Genetic analysis indicated that NPR-15 functions in the same genetic pathway as SEK-1 to regulate high-salt attractive learning. Furthermore, NPR-15 is expressed in AIA interneurons which receive synaptic inputs from ASER, ASH, and ADF chemosensory neurons in which SEK-1 signaling functions for high-salt attractive learning. NLP-3 is a neuropeptide that is abundantly expressed in ASER, ASH, and ADF. The *nlp-3* mutant exhibited a minor yet significant defect in high-salt attractive learning. Furthermore, genetic analysis suggested that SEK-1 and NLP-3 function in the same genetic pathway in high-salt attractive learning, and NPR-15 might be a receptor of NLP-3 in the regulation of high-salt attractive learning. Therefore, based on these findings, we propose that SEK-1 signaling may regulate high-salt attractive learning by controlling the neuropeptide signaling from chemosensory neurons to downstream interneurons such as AIA. As mentioned above, known downstream direct targets of SAPK signaling include some transcription factors and protein kinases, which can provide transcriptional and post-transcriptional regulation in gene expression. NSY-1-SEK-1 p38 signaling promotes the transcription of antimicrobial genes, such as *nlp-29*, in the hypodermis and *tph-1*, the serotonin synthesis gene, in ADF in response to infection (Pujol et al., 2008; Shibers et al., 2009). We examined the possibility that *nlp-3* transcription may be regulated by SEK-1 signaling. However, quantitative PCR analysis did not detect any change in *nlp-3* expression caused by the *sek-1* mutation. SEK-1 signaling might modulate NLP-3 through post-transcriptional regulation, such as neuropeptide processing, which requires further investigation. Meanwhile, it should be noted that the mild effect of the *nlp-3* mutation on salt chemotaxis learning may be a reason for not observing the additive effect on salt chemotaxis learning by the *nlp-3; sek-1* double mutation. Finally, since the high-salt chemotaxis defects of the *nlp-3* and *npr-15* mutants exhibited a less significant effect than those of the *sek-1* mutants, SEK-1 may regulate not only one neuropeptide-neuropeptide receptor signaling but also multiple neuropeptides signaling or different kinds of signal transductions mediated by other neurotransmitters such as monoamines, to mediate salt chemotaxis learning. Further genetic and biochemical research will be required to confirm the hierarchical relationship between SEK-1 signaling, a neuropeptide NLP-3, and a neuropeptide receptor NPR-15.

In conclusion, our study demonstrates that the SAPK signaling pathway is required for salt chemotaxis learning. The MLK-1-MEK-1 JNK signaling pathway promotes avoidance learning, while the NSY-1-SEK-1 p38 signaling pathway promotes both attraction and avoidance learning that leads to migration to high salt environments. Additionally, KGB-1 plays a key role in salt chemotaxis learning, presumably downstream of the p38/JNK signaling pathways. SEK-1 signaling functions in chemosensory neurons, such as ASH, ADF, and ASER, together with GRK-2, to regulate high-salt attractive learning. In addition, the site of action of SEK-1 signaling in avoidance learning appears to differ. Our results suggest that SEK-1 signaling may regulate high-salt attractive learning via neuropeptide signaling between chemosensory neurons and interneurons.

## Data availability

Strains and plasmids are available upon request. The authors affirm that all data necessary for confirming the conclusions of the article are present within the article, figures, and tables.

## Acknowledgments

We thank Dr. Takeshi Ishihara for *sek-1* cDNA and the *nhr-83* and *ocr-2* promoters, Dr. Roger Pocock for the *rgef-1* promoter, and the *Caenorhabditis* Genetics Center and the National Bioresource Project (Japan) for strains. We would like to thank Enago (www.enago.jp) for the English language review.

## Supporting information captions

**Figure S1. Salt Chemotaxis Learning Assay**

Salt chemotaxis learning assay used in this study. (a) Adult worms were transferred to NGM plates with low (25 mM) and high (100 mM) salt concentrations in the absence or presence of food for 6 h, which is called conditioning. After conditioning, the worms were placed in the center of a 9 cm test plate (shown in b) and allowed to crawl for 45 min. (c) To prepare the test plate with a salt gradient, two cylindrical agar blocks with 0 mM NaCl or 150 mM NaCl were placed at points A and B. After 22 h at 20℃, a salt gradient was formed from 35 mM (point A) to 95 mM (point B).

**Figure S2. components of the JNK and p38 MAPK pathways**

All components of the JNK (left) and p38 MAPK pathways (right) identified in *C. elegans.* Although DLK-1, the ortholog of mammalian MAP3K12, and MKK-4, the ortholog of mammalian MAP2K4, are involved in the p38 cascade in *C. elegans*, their ortholog in mammals is involved in the regulation of JNK signaling.

**Figure S3. Mutants of *jkk-1, dlk-1, and mkk-4* exhibit normal salt chemotaxis learning**

Mutants of *jkk-1* (a), a MAPKK in the JNK pathway, *dlk-1* and *mkk-4* (b), a MAPKKK and a MAPKK in the p38 MAPK pathway, do not exhibit defects in salt chemotaxis learning. The bars and error bars represent the mean values and SEM, respectively. ANOVA with Dunnett’s post hoc test (vs wild-type): N = 9 assays.

**Figure S4. The site of action of SEK-1 in low-salt avoidance learning is different.**

Cell-specific rescue experiments of salt chemotaxis learning defects in the *sek-1* mutant. (a) The low-salt avoidance learning defect could be rescued only by *sek-1* expression using the *sra-6* promoter. Most transgenes with successful rescue of the high-salt attractive learning defect failed to rescue the low-salt avoidance defect. (b) Some promoters were used to mimic the expression pattern of the *sra-6* promoter; low-salt avoidance learning could not be significantly restored. The bars and error bars represent the mean values and SEM, respectively. ANOVA with Bonferroni’s post hoc test (Tg- vs Tg+). **P < 0.01. N = 9 assays.

**Figure S5. Comparison of *nlp-3* expression in the wild type and the *sek-1 (km4)* mutant before and after conditioning.**

A qRT-PCR analysis of *nlp-3*, *nlp-14*, and *nlp-47* mRNA levels in control and conditioned (100 mM / Fed) young adult wild-type and *sek-1(km4)* worms. The levels of these mRNAs are normalized to the levels of the *eef-1* mRNA. The bars and error bars represent the mean values and SEM, respectively. Kruskal*–*Wallis and Mann*–*Whitney tests. N = 12 assays.

**Figure S6. Model of interaction between sensory neurons involved in regulating salt chemotaxis learning by SEK-1 signaling and interneurons involved in salt chemotaxis learning.**

ASER sensory neuron sends synaptic input to AIA, AIB, and AIY interneurons; ASH sensory neurons send synaptic input to AIA, AIB, and AIZ interneurons; ADF sensory neurons send synaptic input to AIA, AIY, and AIZ interneurons. ASER, ASH, and ADF collectively connect with interneuron AIA.

**Figure S7. Salt chemotaxis after conditioning with 50 mM NaCl.**

Mutants of *sek-1*, which exhibit high salt chemotaxis defect after high-salt (100 mM) or low-salt (25 mM) learning, also show lower chemotaxis index after conditioning at 50 mM, a medium salt concentration. The bars and error bars represent the mean values and SEM, respectively. ANOVA with Dunnett’s post hoc test (vs wild-type). ***P < 0.001 N = 9 assays.

**Figure S8. Expression pattern of *sek-1::sl2::cfp* driven by the *odr-3* promoter.**

Although the *odr-3* promoter is known to drive expression mainly in AWC, in this study, the *odr-3* promoter abundantly drove expression of *sek-1::sl2::cfp* cDNA in ASH neurons (yellow arrow), slightly but detectably expressed in ADF and AWB neurons. However, no expression was detected in AWC. We suspect that expression sites driven by the *odr-3* promoter may depend on 3’UTR sequences inserted downstream of ORF. The *unc-54* 3’UTR was used in many studies for transgene expression, whereas we used *unc-2* 3’UTR for *sek-1::sl2::cfp* cDNA expression using the *odr-3* promoter.

(Cyan: CFP; Red: DiO)

**Table S1. Strains and transgenic lines**

**Table S2. Promoters used in rescue experiments**

**Table S3. Primers used in Real-Time PCR**

**Table S4. Immobility index of main strains**

